# ShinyLearner: A containerized benchmarking tool for machine-learning classification of tabular data

**DOI:** 10.1101/675181

**Authors:** Stephen R. Piccolo, Terry J. Lee, Erica Suh, Kimball Hill

## Abstract

Classification algorithms assign observations to groups based on patterns in data. The machine-learning community have developed myriad classification algorithms, which are employed in diverse life-science research domains. When applying such algorithms, researchers face the challenge of deciding which algorithm(s) to apply in a given research domain. Algorithm choice can affect classification accuracy dramatically, so it is crucial that researchers optimize these choices based on empirical evidence rather than hearsay or anecdotal experience. In benchmark studies, multiple algorithms are applied to multiple datasets, and the researcher examines overall trends. In addition, the researcher may evaluate multiple hyperparameter combinations for each algorithm and use feature selection to reduce data dimensionality. Although software implementations of classification algorithms are widely available, robust benchmark comparisons are difficult to perform when researchers wish to compare algorithms that span multiple software packages.

Programming interfaces, data formats, and evaluation procedures differ across software packages; and dependency conflicts may arise during installation. To address these challenges, we created ShinyLearner, an open-source project for integrating machine-learning packages into software containers. ShinyLearner provides a uniform interface for performing classification, irrespective of the library that implements each algorithm, thus facilitating benchmark comparisons. In addition, ShinyLearner enables researchers to optimize hyperparameters and select features via nested cross validation; it tracks all nested operations and generates output files that make these steps transparent. ShinyLearner includes a Web interface to help users more easily construct the commands necessary to perform benchmark comparisons. ShinyLearner is freely available at https://github.com/srp33/ShinyLearner.

## Background

Classification falls under the category of supervised learning, a branch of machine learning. When performing classification, researchers seek to assign observations to distinct groups. For example, medical researchers use classification algorithms to identify patterns that predict whether patients have a particular disease, will respond positively to a particular treatment, or will survive a relatively long period of time after diagnosis[1–11]. Applications in molecular biology include annotating DNA sequencing elements, identifying gene structures, and predicting protein secondary structures[12].

Typically, a classification algorithm is “trained” on a dataset that contains samples (observations) from two or more groups, and the algorithm identifies patterns that differ among the groups. If these patterns are reliable indicators of group membership, the algorithm will be able to accurately assign new samples to these groups and thus may be suitable for broader application. Different research applications require different levels of accuracy before classification algorithms are suitable for broader application. However, even small improvements in accuracy can provide large benefits. For example, if an algorithm predicts drug-treatment responses for 1000 patients and attains accuracy levels that are 2% higher than a baseline method, this algorithm would benefit 20 additional patients. Accordingly, a key focus of classification research in the life sciences is to identify generalizable ways to optimize prediction accuracy.

The machine-learning community have developed hundreds of classification algorithms and have incorporated many of these implementations into open-source software packages[13–18]. Each algorithm has different properties, which affect its suitability for particular applications. In addition, most algorithms support hyperparameters, which alter the algorithms’ behavior and can affect the algorithms’ accuracy dramatically. In addition, feature-selection (or feature-ranking) algorithms can be used in complement to classification algorithms, helping to identify combinations of variables that are most predictive of group membership and aiding in data interpretation[19,20]. With this abundance of options to consider, researchers face the challenge of identifying which algorithm(s), hyperparameter combinations, and features are optimal for a particular dataset.

To improve the odds of making successful predictions, researchers should choose algorithms, hyperparameters, and features based on empirical evidence rather than hearsay or anecdotal experience. Prior studies can provide insight into algorithm performance, but few studies evaluate algorithms comprehensively, and performance may vary widely for different types of data. One way to select these options empirically is via nested cross-validation[21]. With this approach, a researcher divides a single dataset into training and validation sets. Within each training set, the researcher divides the data further into training and validation subsets and then evaluates various options using these subsets. The top-performing option(s) are then used when making predictions on the outer validation set. Alternatively, a researcher might perform a benchmark study, applying (non-nested) cross validation to multiple datasets from a given research domain. After testing multiple algorithms, hyperparameters, and/or feature subsets, the researcher can examine overall trends and identify options that tend to perform well[22,23]. With either approach, it is ideal to evaluate a comprehensive set of options. However, several challenges make it difficult to perform such evaluations effectively:

- Researchers may wish to compare algorithms that have been implemented in different software packages. Although many machine-learning packages allow users to execute algorithms programmatically, application programming interfaces (APIs) are not standardized, and they are implemented in diverse programming languages.
- Different software implementations use different techniques for evaluating algorithm performance, so it is difficult to ensure that comparisons are consistent.
- Input and output formats differ by software implementation, thus requiring custom efforts to prepare data and interpret results.
- When installing the software, researchers typically must install a series of software dependencies. Installation requirements often differ by operating system, and versioning conflicts can arise[24].

To reduce these barriers, we created ShinyLearner. For this open-source project, we have integrated existing machine-learning packages into containers, which provide a consistent interface for performing benchmark comparisons of classification algorithms. ShinyLearner can be installed on Linux, Mac, or Windows operating systems, with no need to install software dependencies other than the Docker containerization software. ShinyLearner currently supports 53 classification algorithms and 1300+ hyperparameter combinations across these algorithms; users can perform automatic hyperparameter tuning via nested cross validation. In addition, ShinyLearner supports 16 feature-selection algorithms, enabling researchers to reduce data dimensionality before performing classification (via nested cross validation). New algorithms can be integrated in an extensible manner.

ShinyLearner is designed to be friendly to non-computational scientists—no programming is required. We provide a Web-based tool (http://bioapps.byu.edu/shinylearner) to guide users through the process of creating the Docker commands necessary to execute the software. ShinyLearner supports a variety of input formats and produces output files in “tidy data” format[25], thus making it easy to import results into external tools. Even though other machine-learning packages support nested cross validation, these evaluations may occur in a “black box.” ShinyLearner tracks all nested operations and generates output files that make this process transparent.

Below we describe ShinyLearner in more detail and illustrate its use via benchmark evaluations. We evaluate 10 classification algorithms and 10 feature-selection algorithms on 10 biomedical datasets. In addition, we assess the effects of hyperparameter optimization on predictive performance, provide insights on model interpretability, and consider practical elements of performing benchmark comparisons.

## Methods

ShinyLearner encapsulates open-source, machine-learning packages into Docker images[26], which are available on Docker Hub (https://hub.docker.com/r/srp33/shinylearner/). Currently, ShinyLearner supports algorithms from scikit-learn, Weka, mlr, h2o, and Keras (with a TensorFlow backend)[13–15,27–29]. To facilitate user interaction, to harmonize execution across the tools, and to evaluate predictive performance, ShinyLearner uses shell scripts, Python scripts, R scripts, and Java code[30–32]; these are included in the Docker images. To perform an analysis, the user executes a shell command, specifying arguments to indicate the location(s) of the input files, which algorithms to use, whether to perform Monte Carlo or k-fold cross validation, etc. The analysis is executed within a container, and output files are saved to a directory that the user specifies. TensorFlow provides support for execution on graphical processing units, which requires a slightly different software configuration, so we provide a separate Docker image that enables this feature (https://hub.docker.com/r/srp33/shinylearner_gpu/). All changes to the ShinyLearner code are tested via continuous integration[33]; build status can be viewed at https://travis-ci.org/srp33/ShinyLearner.

Figure S1 shows an example ShinyLearner command that a user might execute. For convenience, and to help users who have limited experience with Docker or the command line, we created a Web-based user interface where users can specify local data paths, choose algorithms from a list, and select other settings (https://bioapps.byu.edu/shinylearner). After the user has made these selections, the Web interface generates a Docker command, which the user can copy and paste; Windows Command Line, Mac Terminal, and Linux Terminal commands are generated. We used the R Shiny framework to build this web application[34].

ShinyLearner interfaces with each third-party machine-learning package via shell scripts wrap that around the software’s API. For each algorithm, one shell script specifies the algorithm’s default hyperparameters. In most cases, additional shell scripts specify alternative hyperparameters. The classification algorithms in ShinyLearner span methodological categories, including linear models, kernel-based techniques, tree-based approaches, Bayesian models, distance-based methods, ensemble approaches, and neural networks. In selecting algorithms to include, we focused primarily on implementations that can handle discrete and continuous data values, support multiple classes, and produce probabilistic predictions. For each algorithm, we reviewed documentation for the third-party software and identified a representative variety of hyperparameter options. Admittedly, these selections are somewhat arbitrary and inexhaustive. However, they can be extended with additional options. We excluded some algorithm implementations and hyperparameter combinations because errors occurred when we attempted to execute them or because they failed to achieve reasonable levels of classification accuracy on simulated data.

Additional algorithms (and hyperparameter combinations) can be incorporated into ShinyLearner. The sole requirements are that they have been implemented as free and open-source software and provide an API (that can be executed via Linux command-line scripts). Users who wish to extend ShinyLearner must:

1. Identify any software dependencies that the new algorithm requires. If those dependencies are not currently included in the ShinyLearner image, the user must modify the ShinyLearner Dockerfiles accordingly.
2. Create bash script(s) that accepts specific arguments and invoke the new algorithm.
3. Request that these changes be included in ShinyLearner via a GitHub pull request.

ShinyLearner supports the following input-data formats: tab-separated value (.tsv), comma-separated value (.csv), and attribute-relation file format (.arff). When tab-separated or comma-separated files are used, column names and row names must be specified; by default, rows must represent samples (observations) and columns must represent features (variables). However, transposed versions of these formats can be used (features as rows and samples as columns); in these cases, the user should use “.ttsv” or “.tcsv” as the file extension. ShinyLearner accepts files that have been compressed with the gzip algorithm (using “.gz” as the file extension). Users may specify more than one data file as input, after which ShinyLearner will identify sample identifiers that overlap among the files and merge on those identifiers. If the user specifies, ShinyLearner will scale numeric values, one-hot encode categorical variables[35], and impute missing values.

ShinyLearner supports two schemes for evaluating predictive performance: Monte Carlo cross validation and k-fold cross validation[36,37]. In Monte Carlo cross validation, the data are split randomly into a training and validation set; the algorithm is allowed to access the class labels for the training data only. Later the algorithm makes predictions for the validation samples, and the accuracy of those predictions is evaluated using various metrics. Typically, this process is repeated many times to derive confidence intervals for the accuracy metrics.

In k-fold cross validation, the process is similar, except that the data are partitioned into evenly sized groups and each group is used as a validation set through rounds of training and testing. When multiple algorithms or hyperparameter combinations are employed, ShinyLearner evaluates nested training and validation sets, with the goal of identifying the optimal combination for each algorithm. Then it uses these selections when making predictions on the outer validation set. Nested cross validation is also used for feature selection; a feature-selection algorithm ranks the features within each nested training set, and different quantities of top-ranked features are used to train the classification algorithm. The feature subsets that perform best are used in making the outer validation-set predictions. Hyperparameter optimization and feature selection may be combined; however, such analyses are highly computationally intensive for large benchmarks.

All outputs are stored in tab-delimited files, thus enabling users to import results directly into external analysis tools. ShinyLearner produces output files that contain the following information for each combination of algorithm, hyperparameters, and cross-validation iteration: 1) predictions for each sample, 2) classification metrics, 3) execution times, and 4) standard output, including a log that indicates the arguments that were used, thus supporting reproducibility. When nested cross-validation is performed, ShinyLearner produces output for every hyperparameter combination that was tested in the nested folds and indicates which combination performed best for each algorithm.

ShinyLearner supports the following classification metrics:

- AUROC (Area under the receiver operating characteristic curve)[38]
- Accuracy (proportion of samples whose discrete prediction was correct)
- Balanced accuracy (to account for class imbalance)
- Brier score[39]
- F1 score[40]
- False discovery rate
- False negative rate
- False positive rate
- Matthews correlation coefficient[41]
- Mean misclassification error
- Negative predictive value
- Positive predictive value
- Recall (sensitivity)
- True negative rate (specificity)
- True positive rate (sensitivity)

To calculate these metrics and to perform other data-processing tasks, ShinyLearner uses the AUC[42], mlr[15], dplyr[43], data.table[44], and readr[45] packages. For multiclass problems, ShinyLearner allows the underlying machine-learning packages to use whatever strategy they have implemented for classifying with multiple classes. ShinyLearner then calculates performance metrics in a one-versus-rest manner and averages results across the class options.

When feature selection is performed, each algorithm produces a ranked list of features for each nested training set. To aid the user in understanding which features are most informative, ShinyLearner aggregates these ranked lists using the *Borda count* method[46]. These aggregate rankings are stored in tab-delimited output files.

### Availability of source code and requirements

- *Project name*: ShinyLearner
- *Project home page*: https://github.com/srp33/ShinyLearner
- *Operating system(s)*: Any operating system on which Docker can be installed
- *Programming languages*: Java, Python, R, bash
- *Other requirements*: Docker (https://docker.com)
- *License*: MIT

The steps of preparing the data and executing ShinyLearner for the results described in this article are in a Jupyter notebook (see https://github.com/srp33/ShinyLearner/blob/master/Demo/Execute_Algorithms.ipynb). The code for creating the figures in this manuscript can be found (and re-executed) in a Code Ocean capsule (https://doi.org/10.24433/CO.5449763.v1). We used the ggplot2 and cowplot packages[47,48] to create figures.

## Analyses

ShinyLearner enables researchers to perform classification benchmark studies. To illustrate this functionality, we performed three types of benchmark: 1) basic classification with default hyperparameters, 2) classification with hyperparameter optimization, and 3) classification with feature selection. For each analysis, we used 10 classification algorithms:

- `keras/dnn` – Deep neural networks (implemented in Keras/TensorFlow)[27,29,49]
- `mlr/h2o.randomForest` – Random forests (implemented in mlr, h2o)[15,28]
- `mlr/mlp` – Multilayer perceptron (mlr)[50]
- `mlr/xgboost` – xgboost (mlr)[51]
- `sklearn/decision_tree` – Decision tree (implemented in scikit-learn)[13,52]
- `sklearn/logistic_regression` – Logistic regression with the LIBLINEAR solver (scikit-learn)[53]
- `sklearn/svm` – Support vector machines (scikit-learn)[54]
- `weka/HoeffdingTree` – Hoeffding tree (implemented in Weka)[14,55]
- `weka/MultilayerPerceptron` – Multilayer perceptron (Weka)
- `weka/SimpleLogistic` – Simple logistic regression (Weka)[56]

In the third analysis, we used 10 feature-selection algorithms:

- `mlr/kruskal.test` – Kruskal-Wallis rank sum test (mlr)[57]
- `mlr/randomForestSRC.rfsrc` – Permuted random forests (mlr)[58]
- `sklearn/mutual_info` – Mutual information (scikit-learn)[59]
- `sklearn/random_forest_rfe` – Random forests—recursive feature elimination (scikit-learn)[60,61]
- `sklearn/svm_rfe` – Support vector machines—recursive feature elimination (scikit-learn)[61]
- `weka/Correlation` – Pearson’s correlation (Weka)[62]
- `weka/GainRatio` – Information gain ratio (Weka)[52]
- `weka/OneR` – OneR (Weka)[63]
- `weka/ReliefF` (Weka)[64]
- `weka/SymmetricalUncertainty` – Symmetrical uncertainty (Weka)[65]

In each analysis, we used 5 rounds of Monte Carlo cross validation. For the second and third analyses, we used 3 rounds of *nested* Monte Carlo cross validation for each *outer* round of cross validation. In the third analysis, we evaluated the top-ranked 1, 3, 5, 10, 15, 20, 50, and 200 features and identified the best of these options via nested cross validation. In evaluating the results, we focused on area under the receiver operating characteristic curve (AUROC) because this metric can be applied to probabilistic predictions and accounts for class imbalance.

As an initial test, we generated a “null” dataset using numpy[66]. We used this dataset to verify that ShinyLearner produces classification results in line with random-chance expectations when no signal is present. This dataset consisted of 20 numeric variables (mean = 0, standard deviation = 1) and 10 categorical variables across 500 simulated samples. AUROC values for all classification algorithms were near 0.5, as expected by random chance, irrespective of whether hyperparameter optimization or feature selection was performed (Figure S2).

Next, we collected 10 biomedical datasets from the Penn Machine Learning Benchmarks repository[67]:

- Acquired Immune Deficiency Syndrome (AIDS) categorical data[68]
- Thyroid disease[52]
- Breast cancer[69]
- Dermatology[70]
- Diabetes
- Hepatitis[71]
- Iris[72]
- Liver disorder[73]
- Molecular biology (promoter gene sequences)[74]
- Yeast[75]

These datasets vary by number of samples (minimum = 51; maximum = 7201) and number of features (min = 5; max = 172). For all datasets, we converted categorical variables to multiple binary variables using one-hot encoding. When executing ShinyLearner, we scaled numeric values using scikit-learn’s RobustScaler, which subtracts the median and scales the data based on the interquartile range[76]; accordingly, this method is robust to outliers. In addition, we used ShinyLearner to impute missing values; this method uses the median for numeric variables and the mode for categorical variables.

### Classification analysis with default hyperparameters

Initially, we applied 10 classification algorithms to 10 biomedical datasets using default hyperparameters. Most algorithms made near-perfect predictions for the Thyroid, Dermatology, and Iris datasets, whereas predictions were less accurate overall for the remaining datasets (Figure 1). The `weka/HoeffdingTree` and `sklearn/decision_tree` algorithms often underperformed relative to the other algorithms (Figure 2). Indeed, for half of the datasets, `weka/HoeffdingTree` performed as poorly or worse than would be expected by random chance. The remaining 8 classification algorithms performed relatively well, but predictive performance varied considerably across the datasets (Figure S3). For example, the AUROC for `mlr/mlp` and `sklearn/logistic_regression` was 0.07 higher than the median on the AIDS dataset; the AUROC for `sklearn/svm` was 0.14 lower than the median.

**Figure 1:**
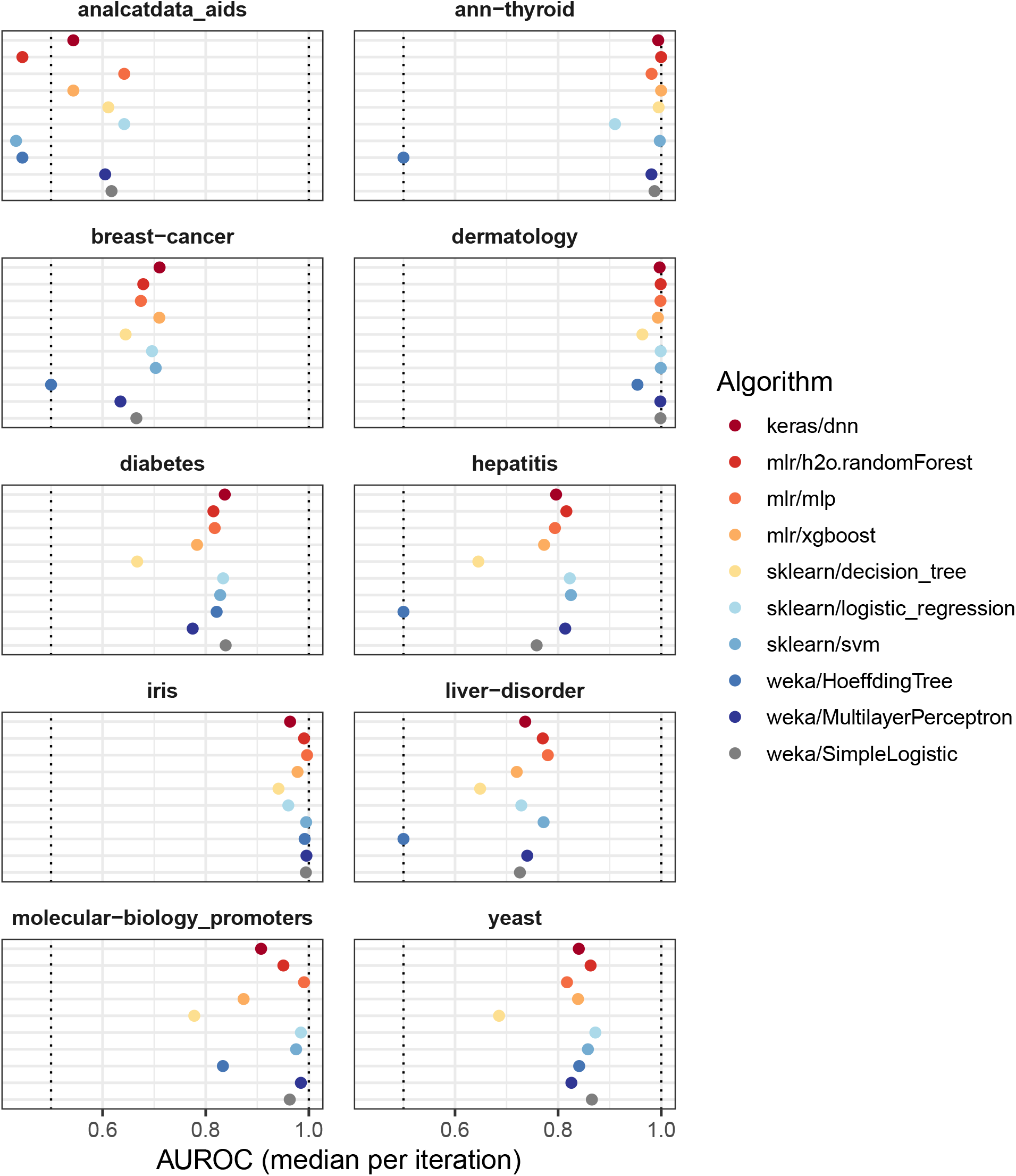
Classification performance per dataset (default hyperparameters). We evaluated the predictive performance of 10 classification algorithms on 10 biomedical datasets. These results were generated using default hyperparameters for each algorithm. We measured predictive performance using the receiver operating characteristic curve (AUROC) and calculated the median across 5 Monte Carlo iterations. Predictive performance differed considerably across and within the datasets.

**Figure 2:**
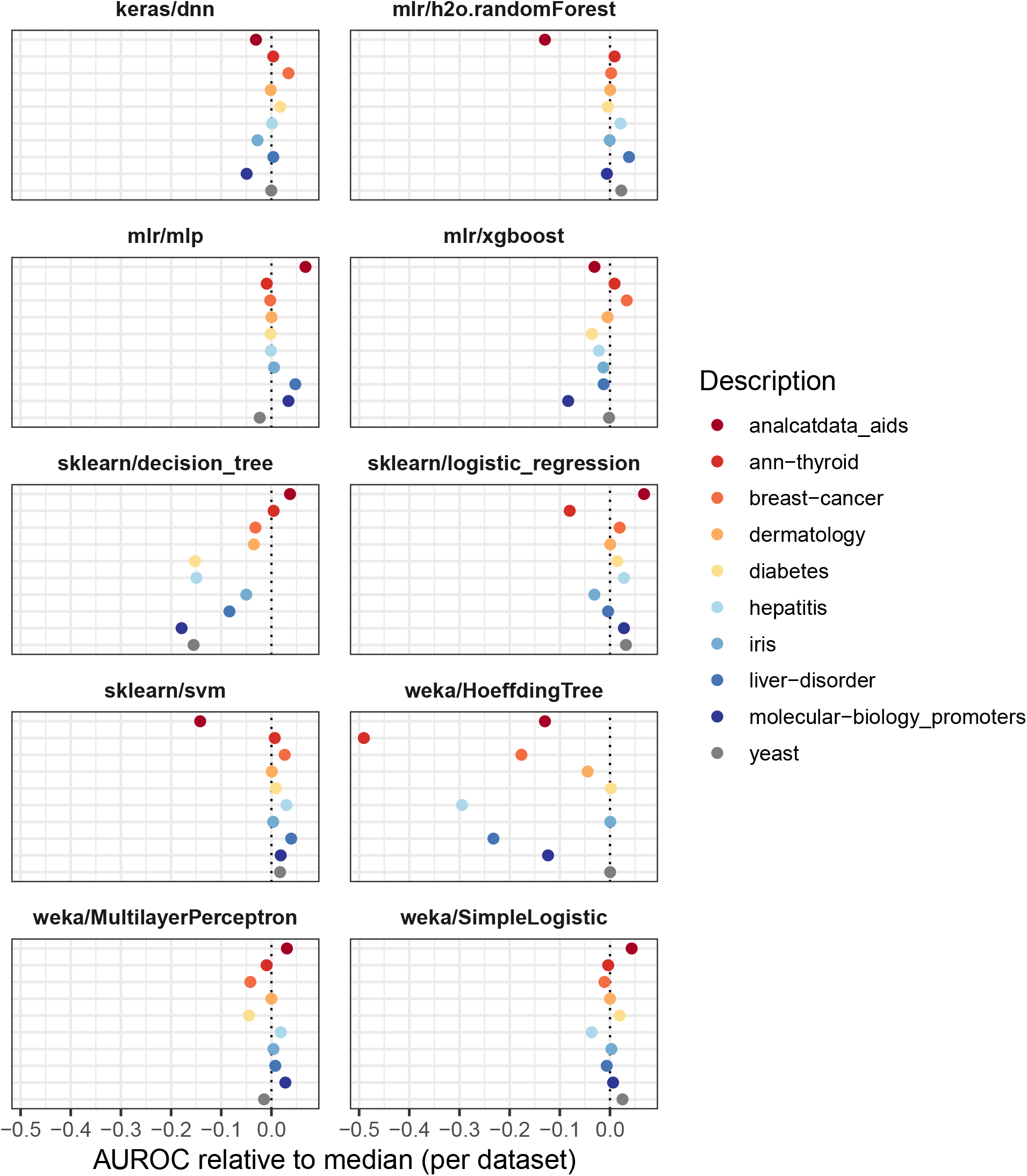
Classification performance per algorithm relative to other classification algorithms (default hyperparameters). We evaluated the predictive performance of 10 classification algorithms on 10 biomedical datasets. These results were generated using default hyperparameters for each algorithm. For each dataset, we calculated the AUROC for each algorithm relative to the median across all algorithms. The `weka/HoeffdingTree` and `sklearn/decision_tree` algorithms underperformed in comparison to the other algorithms.

Across the Monte Carlo iterations for each dataset, the predictive performance of `sklearn/decision_tree` and `weka/MultilayerPerceptron` varied most, whereas `weka/HoeffdingTree` varied least (in part because AUROC was frequently 0.5) (Figure S4). The `keras/dnn` and `mlr/h2o.randomForest` algorithms took longest to execute, whereas `sklearn/svm` and `sklearn/logistic_regression` were among the fastest (and most accurate) algorithms (Figure S5). Two pairs of classification algorithms use similar theoretical approaches but were implemented in different machine-learning libraries; multilayer perceptron was implemented in Weka and mlr; logistic regression was implemented in Weka and scikit-learn. The AUROC values were strongly—but not perfectly—correlated between these pairs of implementations (Figures S6 and S7).

With the exception of `sklearn/decision_tree`, all classification algorithms produced sample-wise, probabilistic predictions. We examined these predictions for the Diabetes dataset and found that the range and shape of these predictions differed widely across the algorithms (Figure 3). Although many classification metrics, including AUROC, can cope with distributional differences, these differences must be considered in multiple classifier systems[77].

**Figure 3:**
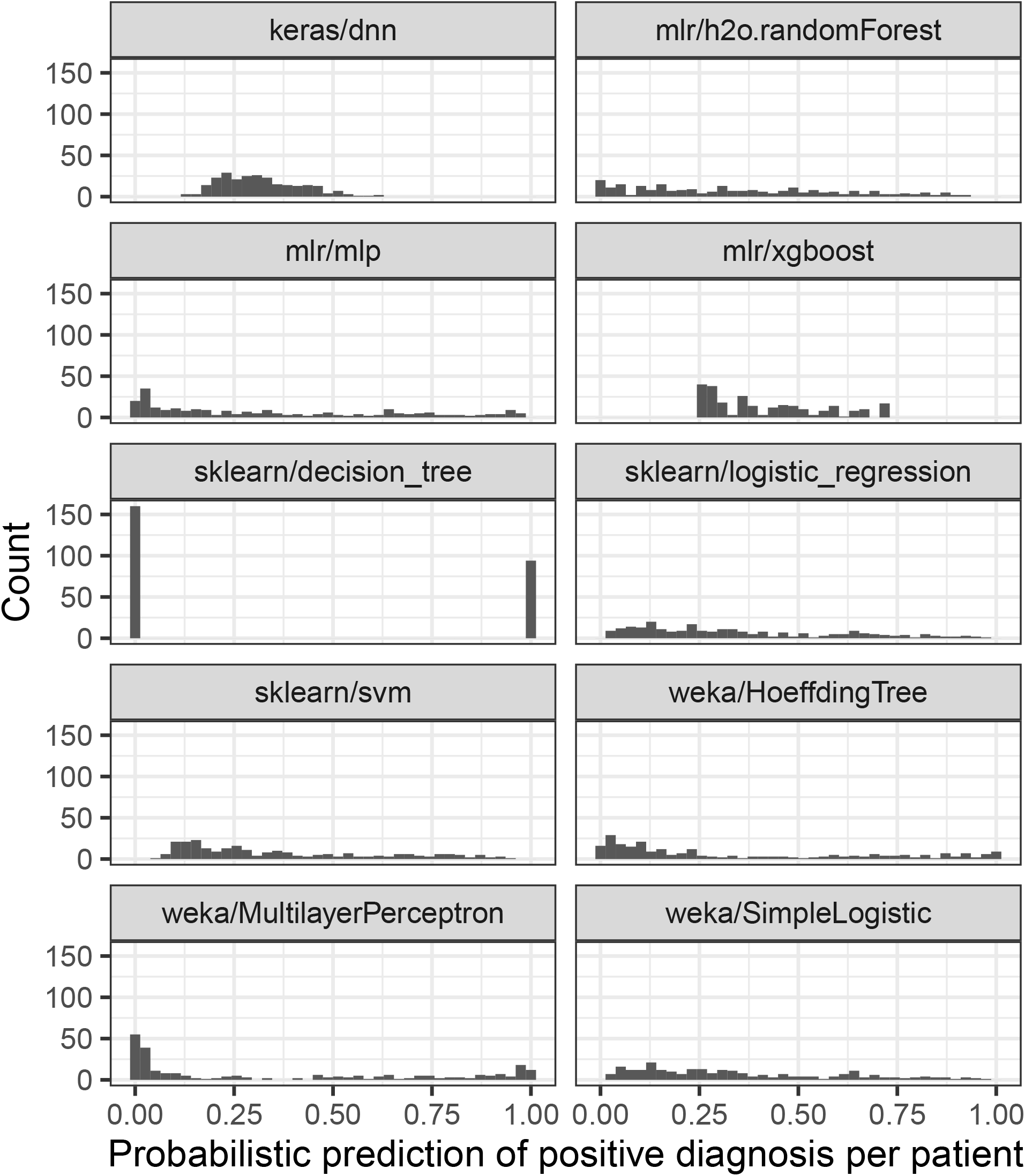
Sample-level predictions for each algorithm on the Diabetes dataset (default hyperparameters). The Diabetes dataset includes a class variable indicating whether or not patients received a positive diagnosis. Each panel of this figure shows positive-diagnosis predictions for each classification algorithm. All algorithms except `sklearn/decision_tree` produced probabilistic predictions. The range and distribution of these predictions differed greatly across the algorithms.

### Classification analysis with hyperparameter optimization

In the second analysis, we applied the same classification algorithms to the same datasets but allowed ShinyLearner to perform hyperparameter optimization via nested cross validation. As few as 2 (`mlr/xgboost`) and as many as 95 (`sklearn/decision_tree` and `weka/MultilayerPerceptron`) hyperparameter combinations were available for each algorithm. In nearly every example, classification performance improved after hyperparameter optimization (Figure 4), sometimes dramatically. The performance improvements were most drastic for the `weka/HoeffdingTree` and `sklearn/decision_tree` algorithms, which often performed poorly with default parameters.

**Figure 4:**
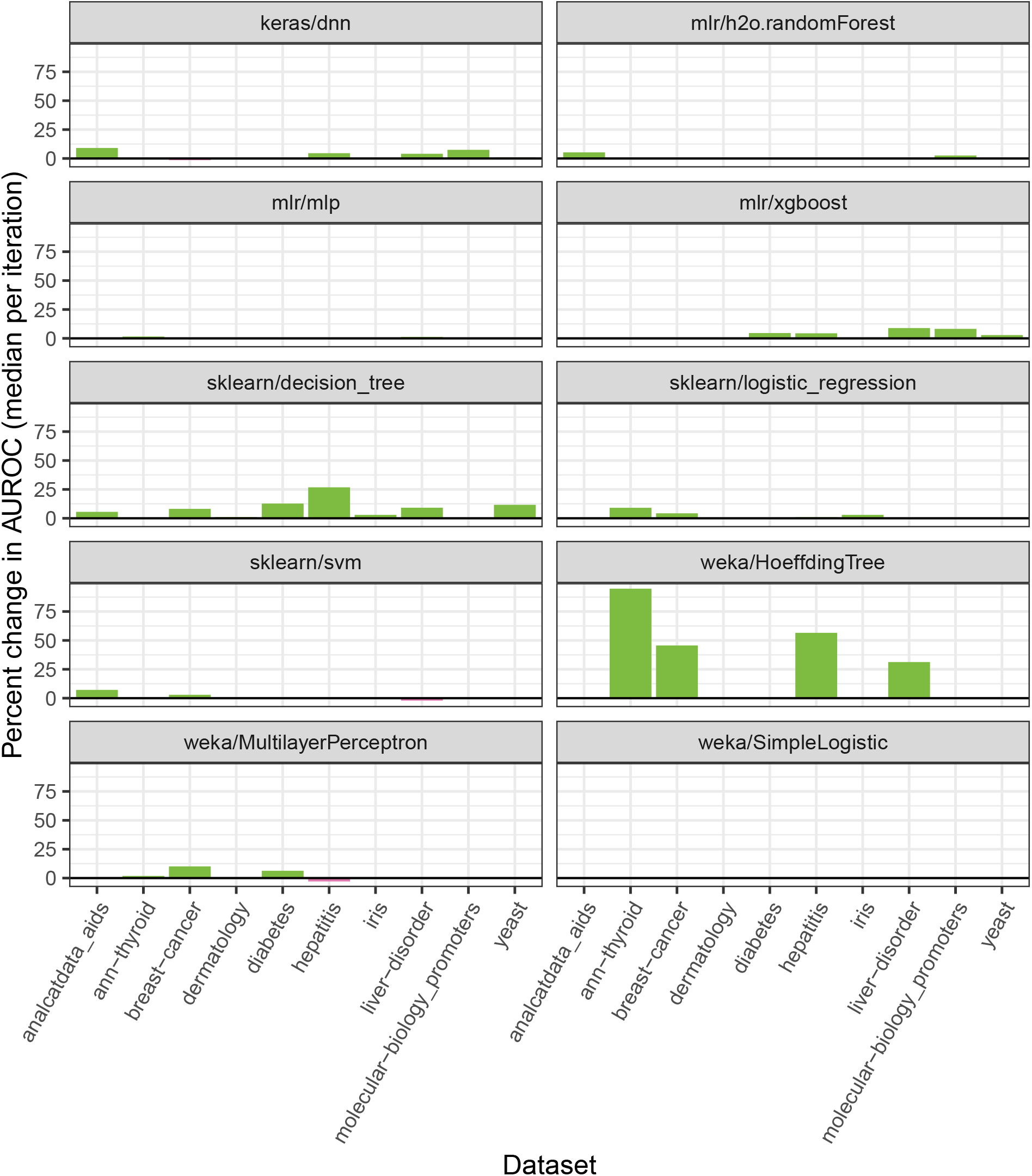
Classification performance when optimizing vs. not optimizing hyperparameters. We tested 10 classification algorithms on 10 biomedical datasets and used nested cross validation to select hyperparameters. To evaluate for change in predictive performance, we calculated the percent change in the median AUROC values when using optimized vs. default hyperparameters. Most algorithms demonstrated improved classification performance with optimized hyperparameters.

ShinyLearner supports 53 hyperparameter combinations for the `keras/dnn` algorithm. Each of these combinations altered the algorithm’s performance at least to a small degree on every dataset (Figure S8). The Thyroid dataset varied least across the hyperparameter combinations, perhaps because the number of instances (n = 7200) was nearly 10 times larger than any other dataset. Generally, this algorithm performed better with a wider architecture containing only two layers. Having a wider structure greatly increases the parameter space of the network and allows it to learn more complex relationships among features, while limiting the network to only two layers prevents overfitting, a common problem when applying neural networks to datasets with a limited number of instances. In addition, adding dropout and L2 regularization also helps to prevent the network from overfitting. In tuning these hyperparameters, we found that a smaller dropout rate, more training epochs, and a smaller regularization rate resulted in higher AUROC values (Figure S9). Figure S10 illustrates for the Diabetes dataset that diagnosis predictions can differ considerably, depending on which hyperparameter combination is used.

### Classification analysis with feature selection

In any dataset, some features are likely to be more informative than other features. We used ShinyLearner to perform feature selection (via nested cross validation) before classification. In total, we evaluated 100 unique combinations of feature-selection algorithm and classification algorithm (with default hyperparameters). In 44% of cases, feature selection increased the median AUROC, whereas it decreased AUROC in 39% of cases (Figure 5). Feature selection sometimes improved the performance of `weka/HoeffdingTree` and `sklearn/decision_tree`, which were the lowest performers without feature selection.

**Figure 5:**
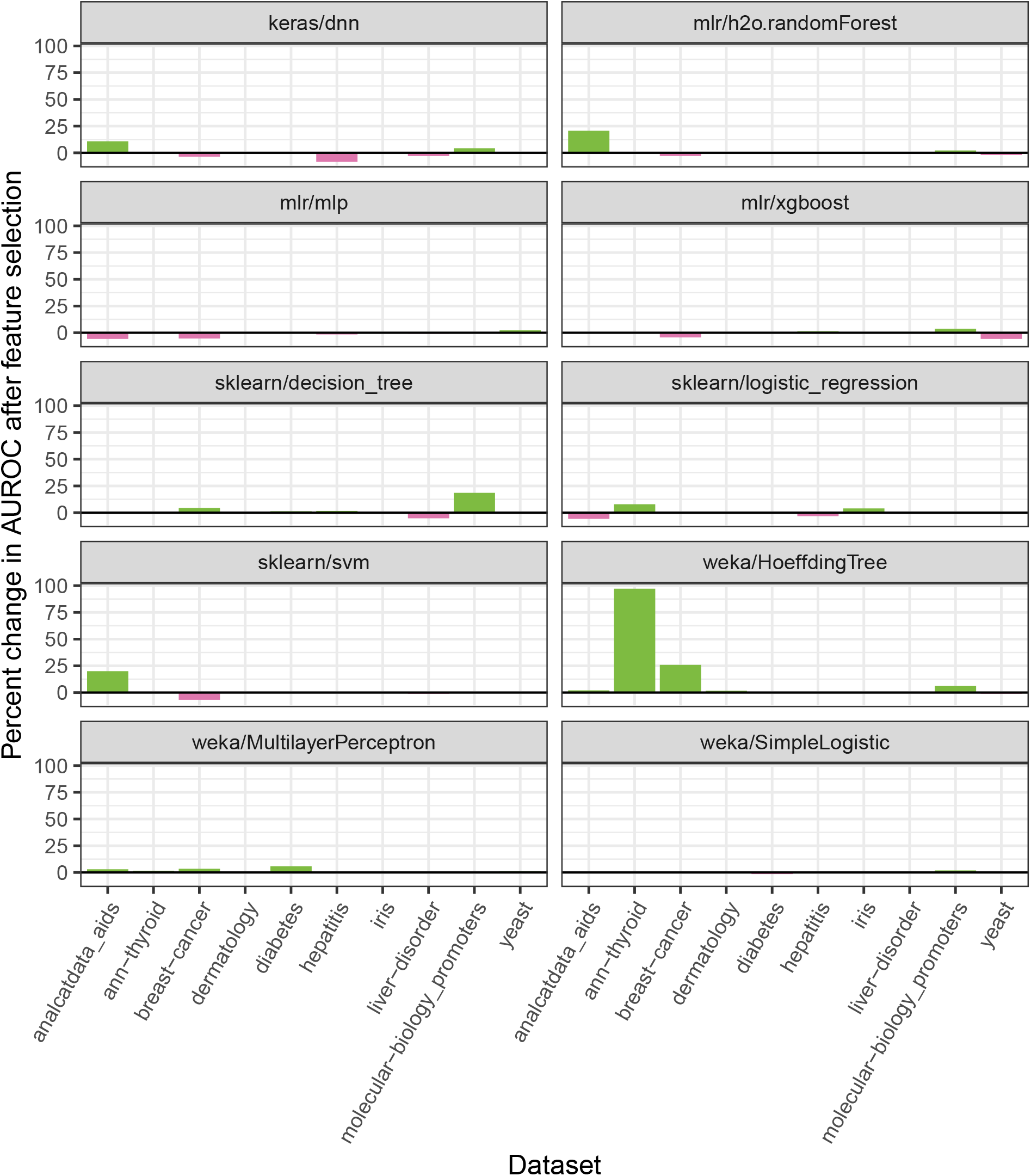
Classification performance when performing feature selection vs. not performing feature selection. In combination with classification, we performed feature selection via nested cross validation on 10 biomedical datasets. For each algorithm, we used default hyperparameters. These plots show the percent change in the median AUROC when using vs. not using feature selection. Although the effects of feature selection varied across the algorithms, median AUROCs increased in many cases.

Figure 6 illustrates the relative predictive ability of each combination of feature-selection and classification algorithms. The `mlr/randomForestSRC.rfsrc` and `sklearn/random_forest_rfe` algorithms performed best on average; both approaches use the Random Forests algorithm to evaluate feature relevance. The `weka/OneR` algorithm, which evaluates a single feature at a time in isolation, performed worst. Across the datasets, the combination of `mlr/randomForestSRC.rfsrc` (feature selection) and `mlr/xgboost` (classification) performed best. Perhaps surprisingly, the combination of `sklearn/svm_rfe` (feature selection) and `sklearn/svm` (classification), which are both based on Support Vector Machines, was ranked in the bottom quartile.

**Figure 6:**
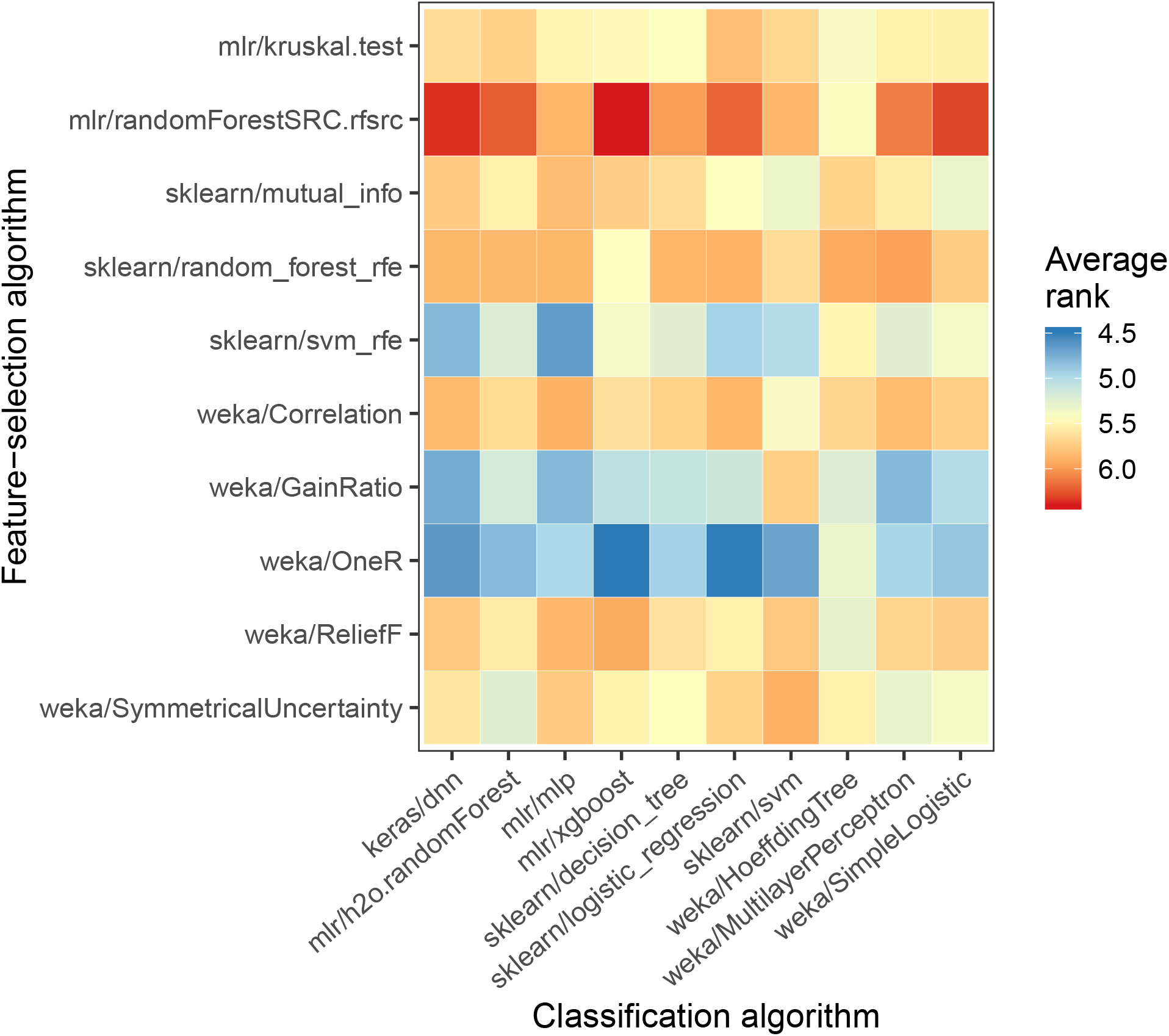
Performance for each combination of classification and feature-selection algorithm. This figure shows classification results for the nested cross-validation folds across each combination of feature-selection algorithm and classification algorithm. Averaged across all datasets and classification algorithms, we ranked the feature-selection algorithms based on AUROC values attained for nested validation sets. For simplicity and consistency across the datasets, this figure shows only the results when the top-5 features were used. Higher average ranks indicate better classification performance.

In seeking to identify the most informative features, ShinyLearner evaluated various quantities of top-ranked features via nested cross validation. Figure 7 illustrates the relative performance of each of these quantities on each dataset. In all cases but one, using one feature performed worst. Generally, a larger number of features resulted in higher AUROC values. However, more features sometimes decreased performance. For example, on the breast-cancer dataset, the highest AUROC values were attained using 3 out of 14 features.

**Figure 7:**
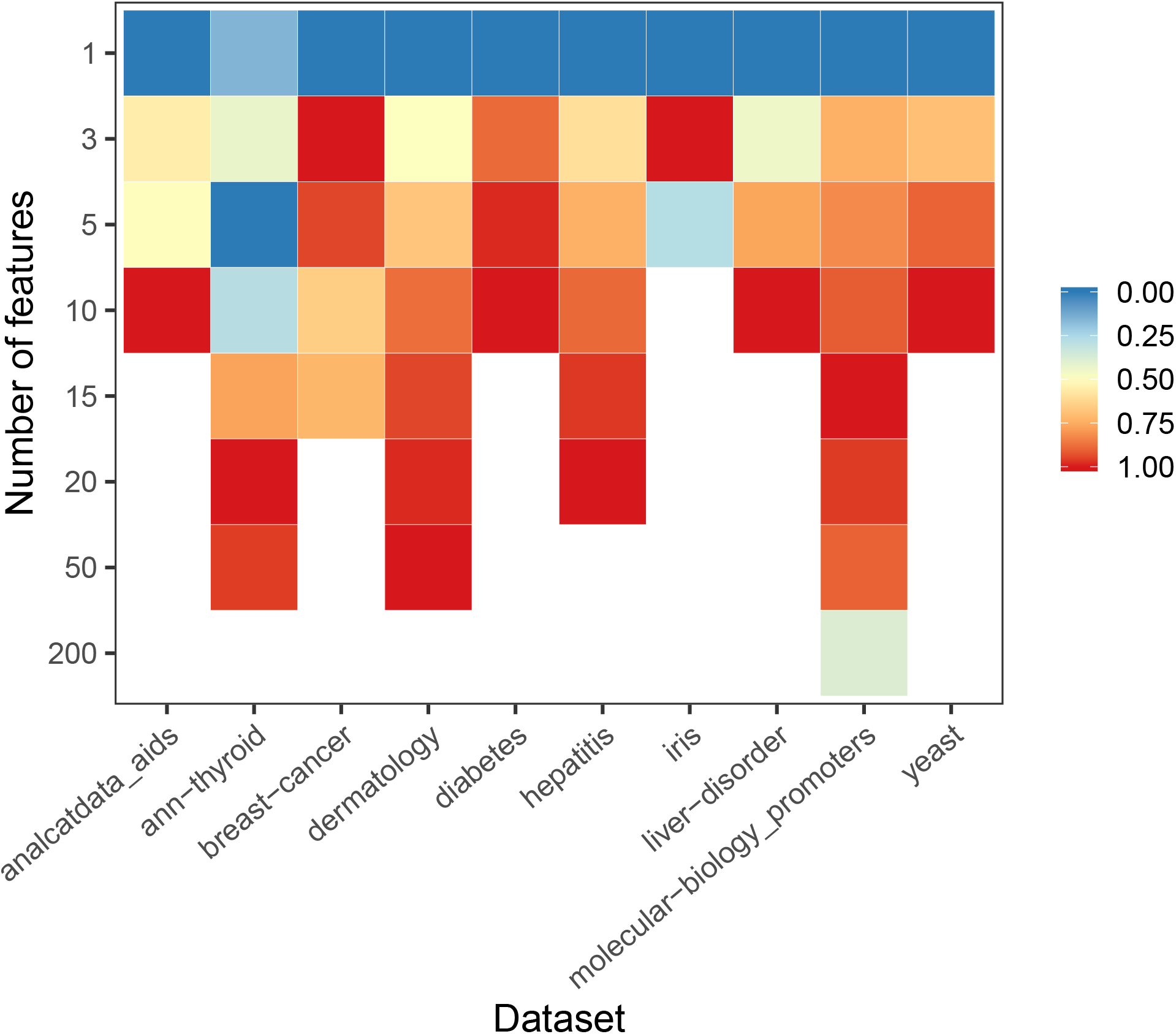
Median classification performance of feature-selection algorithms by number of features. We applied feature selection to each dataset and selected the top *x* number of features. This figure shows which values of *x* resulted in the highest AUROC values for each dataset, averaged across all feature-selection algorithms. Different datasets had different quantities of features; this graph only shows results for *x* values relevant to each dataset. Accordingly, we scaled the AUROC values in each column between zero and one to ensure that the comparisons were consistent across all datasets. Higher values indicate better classification performance. Generally, a larger number of features resulted in better classification performance, but this varied across the datasets.

ShinyLearner can inform users about which features are most informative for classification. In the Dermatology dataset, these feature ranks were highly consistent across the feature-selection algorithms (Figure S11). The goal of this classification problem was to predict a patient’s type of Eryhemato-Squamous disease. Elongation and clubbing of the rete ridges as well as thinning of the suprapapillary epidermis were most highly informative of disease type, whereas features such as the patient’s age were less informative.

## Discussion

The machine-learning community has developed an abundance of algorithms and software implementations of those algorithms. Life scientists use these resources for many research applications. But they face the challenge of identifying which algorithms and hyperparameters will be most accurate and which features are most informative for a given dataset. Many researchers limit classification analyses to a single algorithm, perhaps one that is familiar to them or that has been reported in the literature for a similar study. Others may try a large number of algorithms; however, performing benchmark comparisons in an *ad hoc* manner requires a considerable coding effort and can introduce biases if done improperly. Alternatively, some researchers may develop new algorithms without providing evidence that these algorithms outperform existing ones. We developed ShinyLearner as a way to simplify the process of performing classification benchmark studies.

ShinyLearner does not implement any classification or feature-selection algorithm; rather, it serves as a wrapper around existing software implementations. Currently, algorithms from Weka, scikit-learn, mlr, h2o, and Keras are supported in ShinyLearner. In aggregate, these algorithms represent a diverse range of methodological approaches and thus can support comprehensive benchmark evaluations. On their own, each of the third-party tools encapsulated within ShinyLearner provides a way to optimize hyperparameters programmatically and perform feature selection. In addition, tools such as caret[17], KNIME[18], and Orange[78] provide these options. Thus, in situations where a researcher has programming expertise and is satisfied with the algorithms and tuning functionality available in one of those tools, the researcher might prefer to use these tools directly rather than use ShinyLearner. ShinyLearner is most useful when a researcher:

1. wishes to compare algorithms that have been implemented in multiple machine-learning packages,
2. does not have programming expertise,
3. desires to perform complex operations via nested cross validation, such as evaluating different sizes of feature subsets,
4. wishes to analyze algorithm performance using a tool or programming language that is different than was used to perform classification,
5. wishes to gain deeper insight into decisions made during nested cross validation, and/or
6. seeks to evaluate the tradeoff between predictive accuracy and time of execution.

ShinyLearner is limited to datasets that fit into computer memory. For larger datasets, frameworks such as Apache SystemML support distributed algorithm execution[79]; however, the number of algorithms implemented in these frameworks is still relatively small.

The current release of ShinyLearner supports diverse classification algorithms and hyperparameter combinations; however, this collection is far from exhaustive. Using ShinyLearner’s extensible architecture, the research community can integrate additional algorithms and hyperparameter combinations. In addition, algorithm designers can use our framework to compare their algorithms against competing methods and disseminate their algorithms to the research community.

Containers provide many advantages for software deployment. Tool installation and computational reproducibility are easier because all software components are encapsulated within the container, and container images can be archived and versioned[80]. One other benefit may be less apparent: containerization facilitates the use of diverse programming languages. Distinct components of ShinyLearner are implemented in 4 different programming languages. We chose this approach because we determined that each language was suited to specific types of tasks. We posit that the future of bioinformatics development will increasingly follow this pattern. Furthermore, we advocate for the approach of providing a graphical user interface, such as the Web-based tool we provided for ShinyLearner. Such tools make it easier for users—especially those who have limited command-line experience—to formulate Docker commands.

Our analysis of 10 biomedical datasets, 10 classification algorithms, and 10 feature-selection algorithms confirmed that the choice of algorithm and hyperparameters has a considerable impact on classification performance and selected features. Although some algorithms typically performed better than others, no single algorithm consistently outperformed any other. This finding supports the “No Free Lunch” theorem[81] and confirms that multiple classifier systems hold promise for aggregating evidence across algorithms[82]. Also importantly, algorithm performance is likely to differ according to data characteristics. Algorithms that perform well on “wide” datasets (many features, few samples) may not perform as well on “tall” datasets. Algorithms that perform well with numeric data may not perform as well on categorical or mixed data. These differences highlight the importance of domain-specific benchmark comparisons.

## Declarations

### List of abbreviations

AUROC: Area under receiver operating characteristic curve
API: application programming interface

### Ethics approval and consent to participate

Not applicable.

### Consent for publication

Not applicable.

### Competing interests

The authors declare that they have no competing interests.

### Funding

SRP was supported by internal funds from Brigham Young University. TJL and KH were supported by fellowships from the Simmons Center for Cancer Research at Brigham Young University.

### Author’s contributions

SRP, TJL, and KH helped to develop the software. SRP conceived of the software design with critical input from TJL and KH. ES and SRP performed the analyses described in the manuscript. All authors helped to write the manuscript.

## Notes

https://github.com/srp33/ShinyLearner

https://doi.org/10.24433/CO.5449763.v1

