## Supplementary Material for "ShinyLearner: A containerized benchmarking tool for machine-learning classification of tabular data"

Stephen R. Piccolo[1,*], Terry J. Lee[1], Erica Suh[1], Kimball Hill[1]

1 - Department of Biology, Brigham Young University, Provo, UT, USA

```
mkdir -p "/home/user/OutputData"

docker run --rm -i \
  -v "/home/user/InputData":"/InputData" \
  -v "/home/user/OutputData":"/OutputData" \
  --user $(id -u):$(id -g) \
  srp33/shinylearner:version511 \
  /UserScripts/nestedclassification_montecarlo \
    --data "data.tsv" \
    --description "My_Analysis_Description" \
    --outer-iterations 10 \
    --inner-iterations 5 \
    --classif-algo "/AlgorithmScripts/Classification/tsv/keras/dnn/*" \
    --classif-algo "/AlgorithmScripts/Classification/tsv/mlr/h2o.randomForest/*" \
    --classif-algo "/AlgorithmScripts/Classification/tsv/mlr/xgboost/*" \
    --classif-algo "/AlgorithmScripts/Classification/tsv/sklearn/svm/*" \
    --seed 1 \
    --ohe true \
    --scale true \
    --impute false
```

**Figure S1: Example ShinyLearner command for performing a benchmark comparison.** In this example, the user wishes to place output files in a directory located at `/home/user/OutputData`. To avoid problems with file permissions, this directory should be created before Docker is executed. The `docker run` command builds a container and maps input and output directories from the host operating system to locations within the container (separated by colons). The `--user` directive indicates that the container should execute using the executing user's permissions. The name of the Docker image and tag name are specified (`srp33/shinylearner:version511`) as well as the name of a ShinyLearner script that performs nested, Monte Carlo cross validation (`/UserScripts/nestedclassification_montecarlo`). The remaining arguments indicate the name of the input data file, a description of the analysis, the number of Monte Carlo iterations, the classification algorithms, etc. ShinyLearner provides documentation on each of these arguments as well as a Web application for building such commands dynamically.

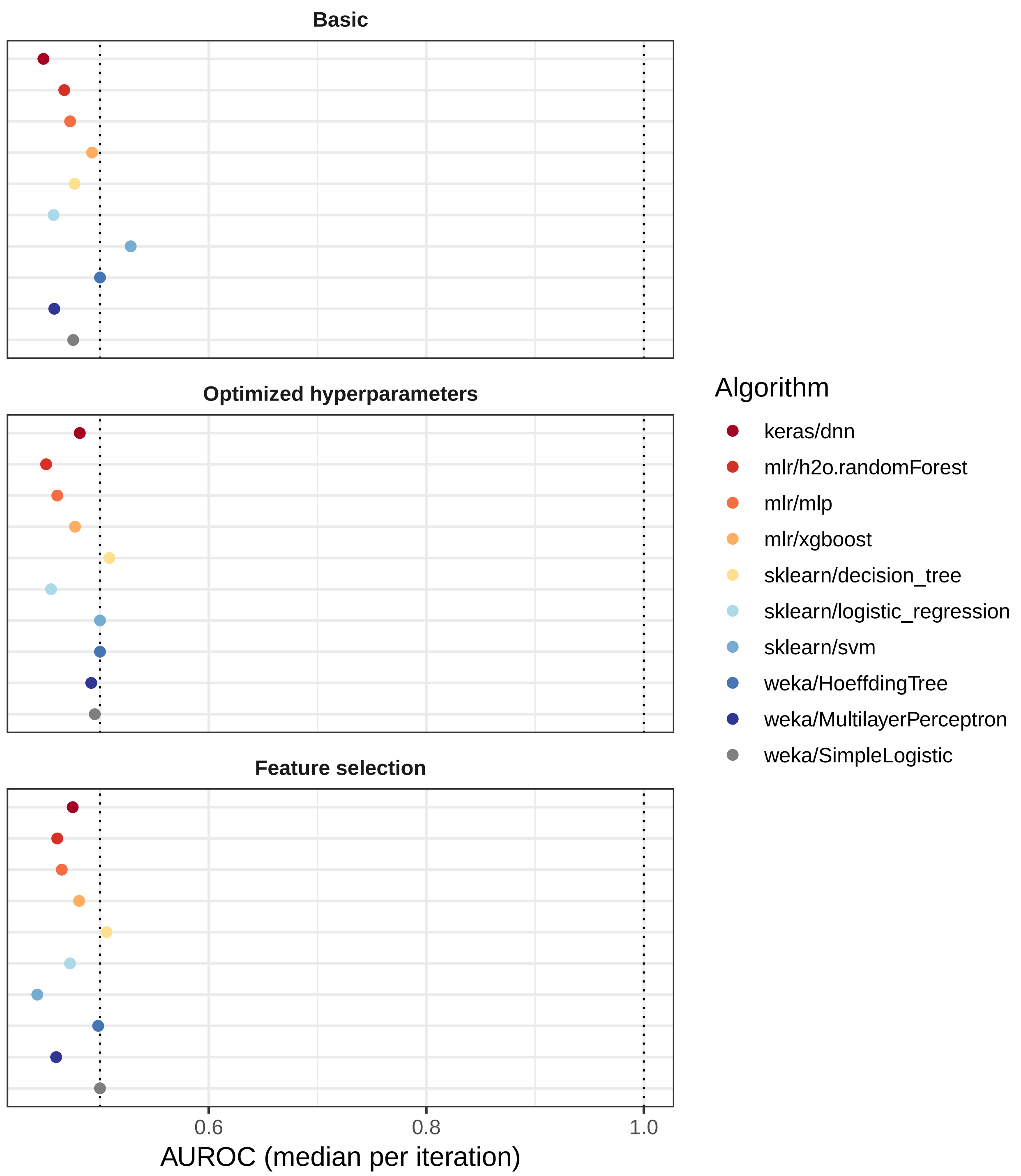


**Figure S2: Classification performance on the "null" dataset.** To verify ShinyLearner's functionality, we randomly generated a "null" dataset and applied 3 types of analysis to the data. In the Basic analysis, default hyperparameters were used for each algorithm. In the second analysis, we used the same algorithms but used nested cross validation to select hyperparameters. In the third analysis, we performed feature selection via nested cross validation. For each analysis, area under the receiver operating characteristic curve (AUROC) was consistently close to 0.5, as expected by random chance.

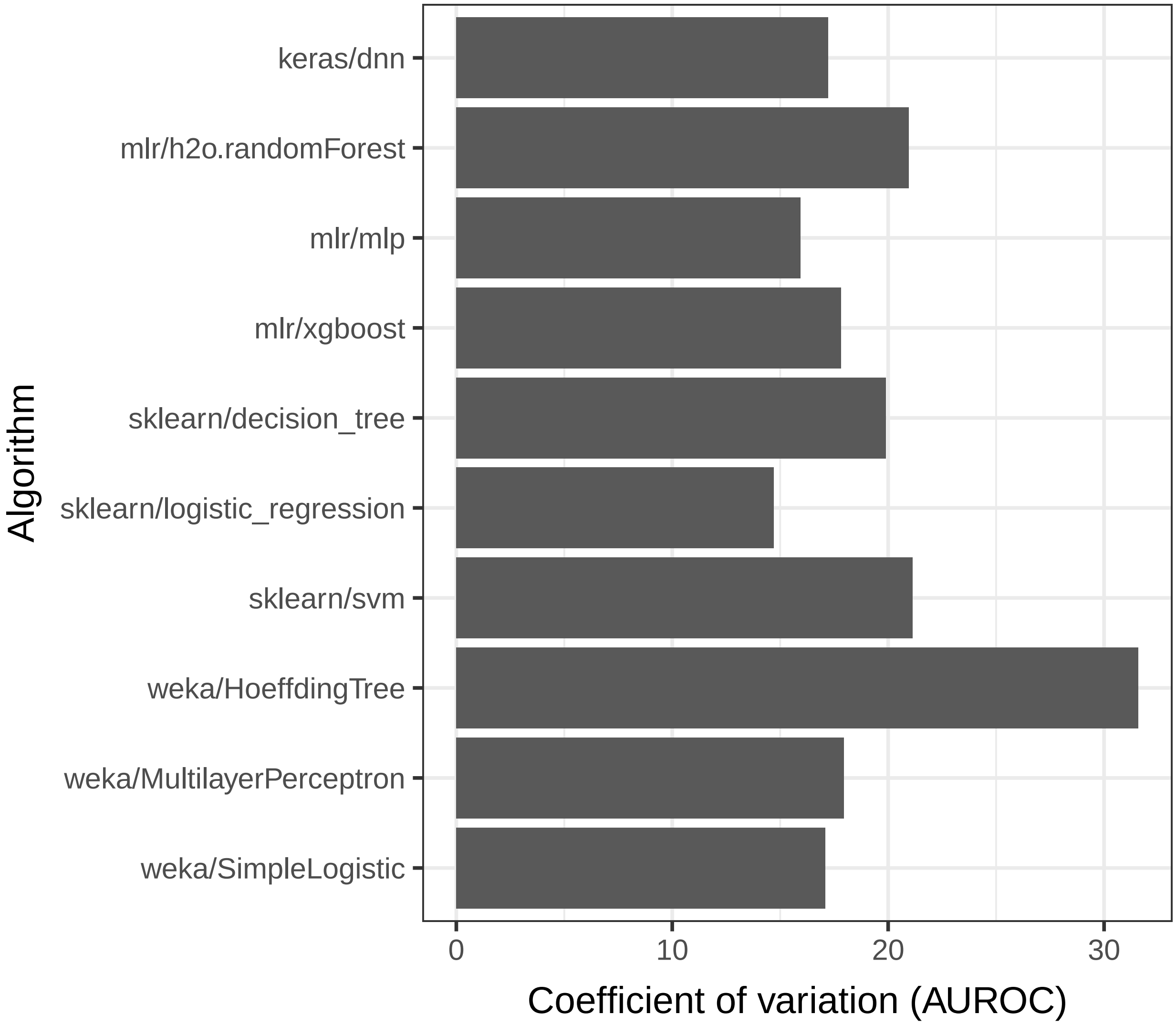


**Figure S3: Consistency of results across datasets for each algorithm (default hyperparameters).** We evaluated the consistency of area under the receiver operating characteristic curve (AUROC) values for each algorithm across the datasets. After calculating the median AUROC across Monte Carlo iterations, we calculated the coefficient of variation across the datasets. The `weka/HoeffdingTree` algorithm varied most across the datasets, while `sklearn/logistic_regression` varied least.

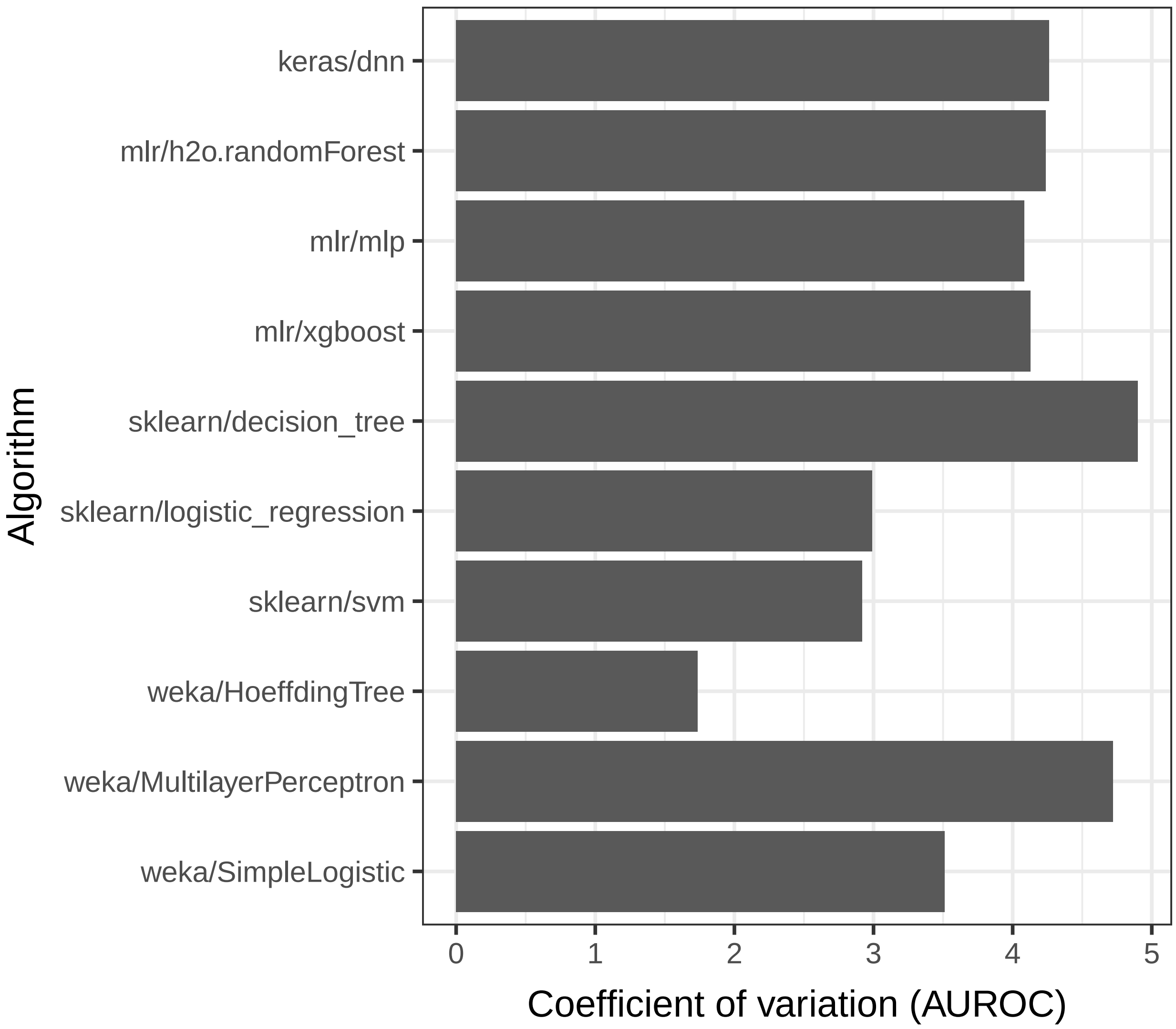


**Figure S4: Consistency of results across Monte Carlo iterations for each algorithm (default hyperparameters).** We evaluated the consistency of area under the receiver operating characteristic curve (AUROC) values across Monte Carlo iterations within each dataset and then calculated the median across the datasets for each algorithm. The `weka/HoeffdingTree` algorithm varied least, while `sklearn/decision_tree` varied most.

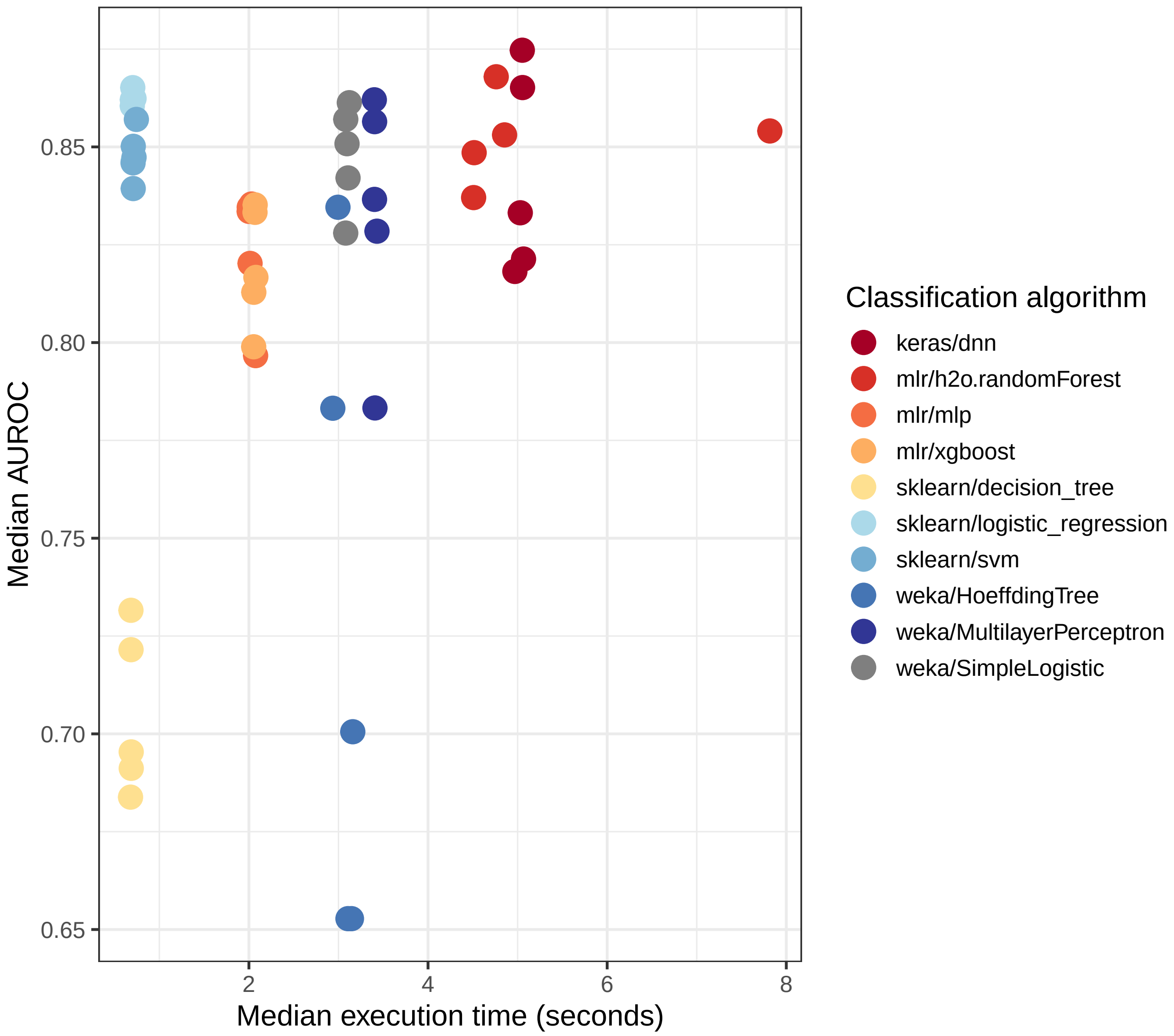


**Figure S5: Relationship between execution time and predictive performance per classification algorithm.** Across 10 biomedical datasets, execution time and area under the receiver operating characteristic curve (AUROC) differed considerably. Each point represents the median value across all datasets for a single Monte Carlo iteration. We observed little to no association between execution time and predictive performance. Some of the best-performing algorithms were also quite fast.

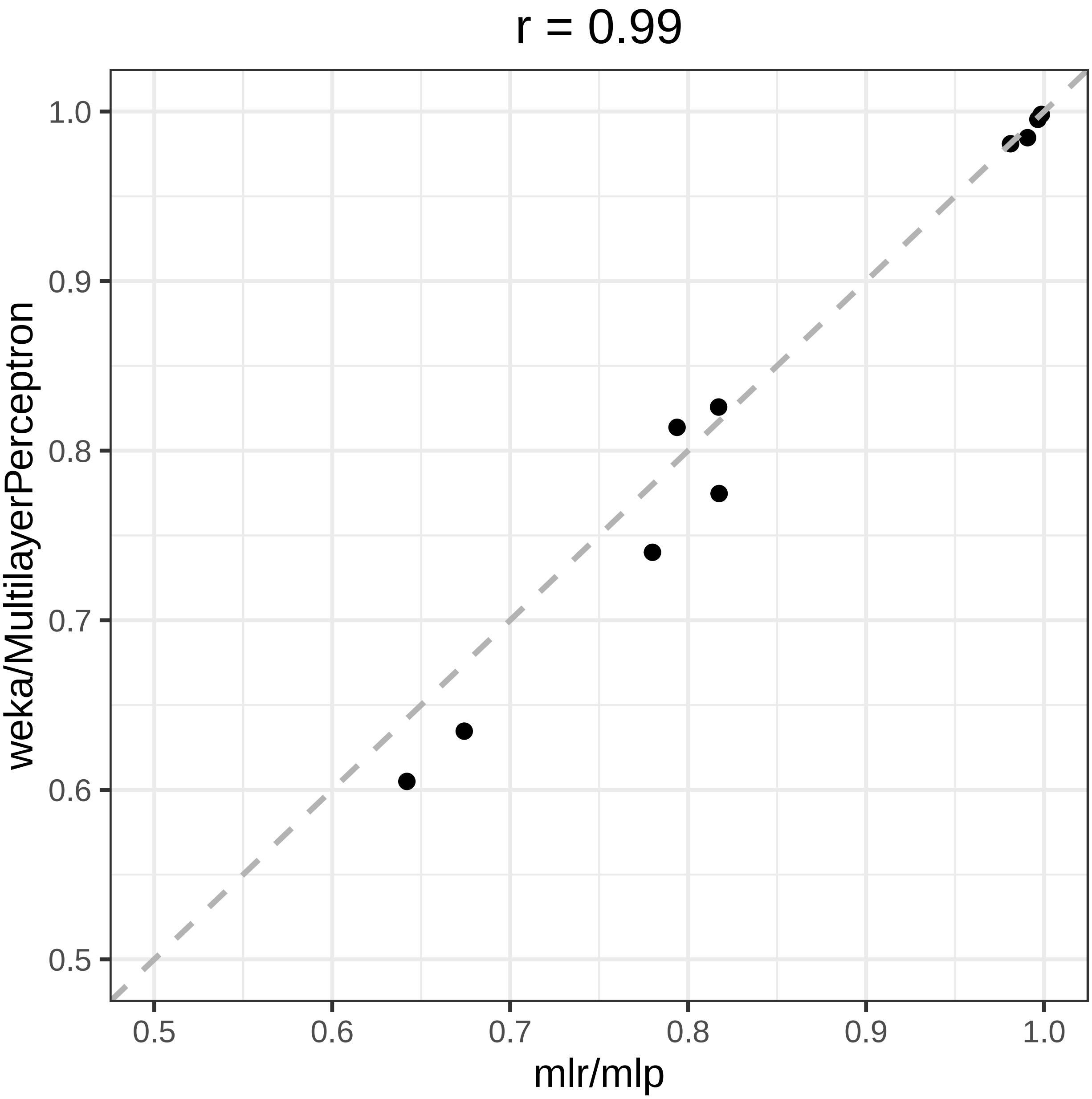


**Figure S6: Comparison of classification performance between two implementations of the multilayer perceptron algorithm (default hyperparameters).** We evaluated the predictive performance (area under the receiver operating characteristic curve) for two implementations of the multilayer perceptron classification algorithm. We compared implementations from the weka and mlr software packages. Predictive performance was highly consistent but not identical. We used Pearson's method to calculate the correlation coefficient.

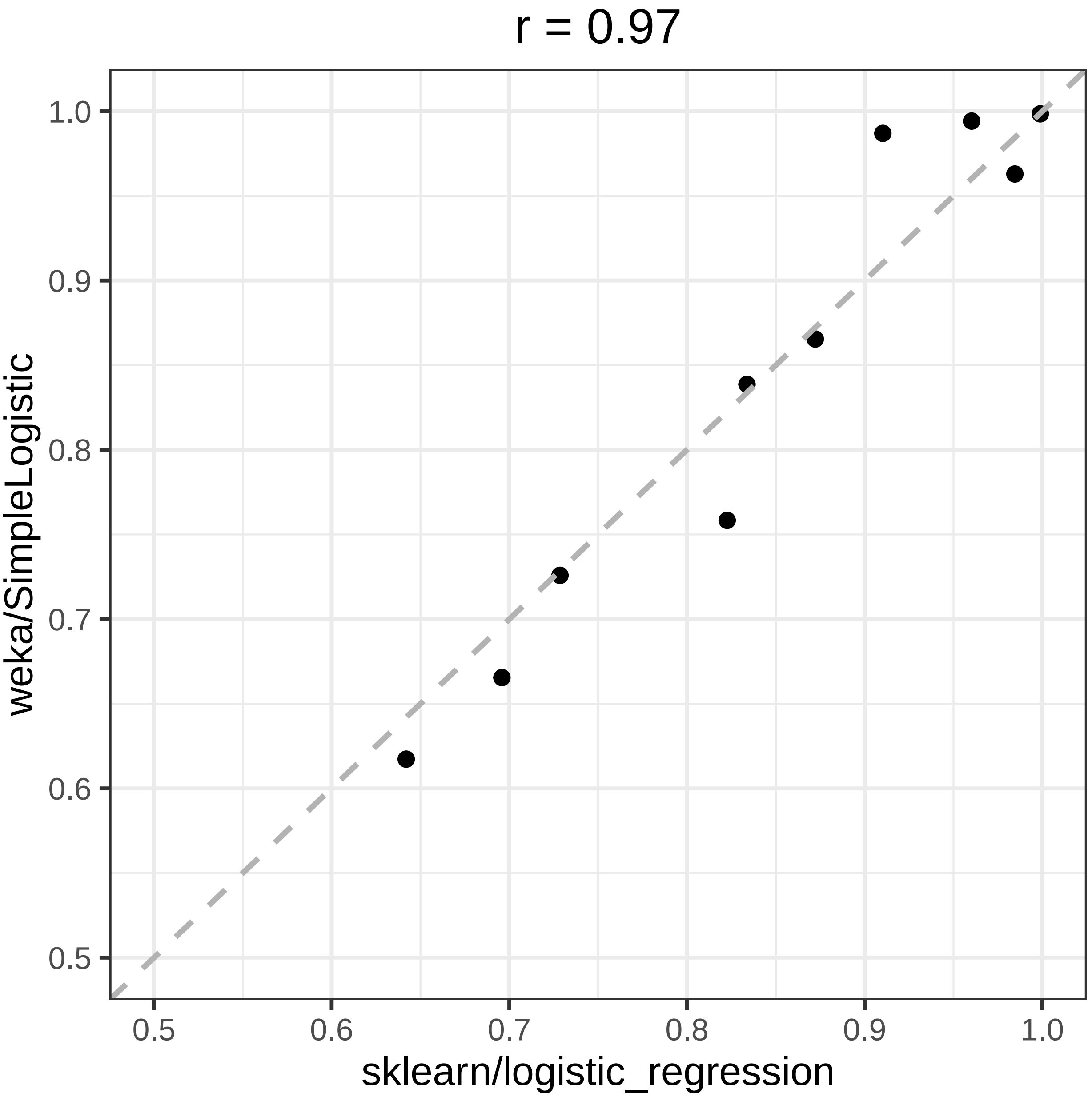


**Figure S7: Comparison of classification performance between two implementations of the logistic regression algorithm (default hyperparameters).** We evaluated the predictive performance (area under the receiver operating characteristic curve) for two implementations of the logistic regression classification algorithm. We compared implementations from the weka and scikit-learn (sklearn) software packages. Predictive performance was highly consistent but not identical. We used Pearson's method to calculate the correlation coefficient.

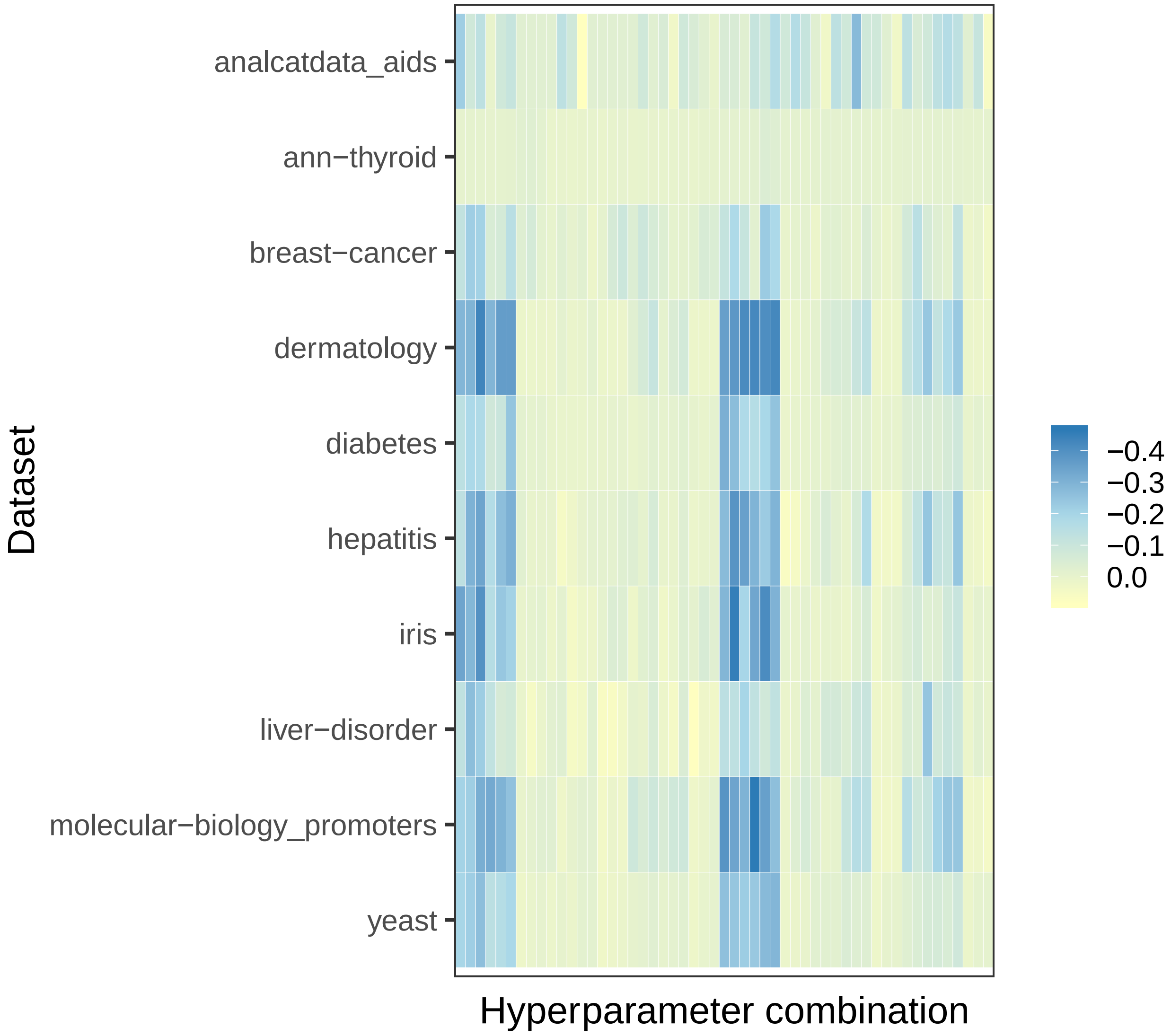


**Figure S8: Performance of different hyperparameter combinations for the keras/dnn classification algorithm.** We evaluated predictive performance for the `keras/dnn` algorithm using 53 different hyperparameter combinations. Each dataset was affected by the combinations to some degree. The Thyroid dataset demonstrated the least variability, possibly due to its large number of instances.

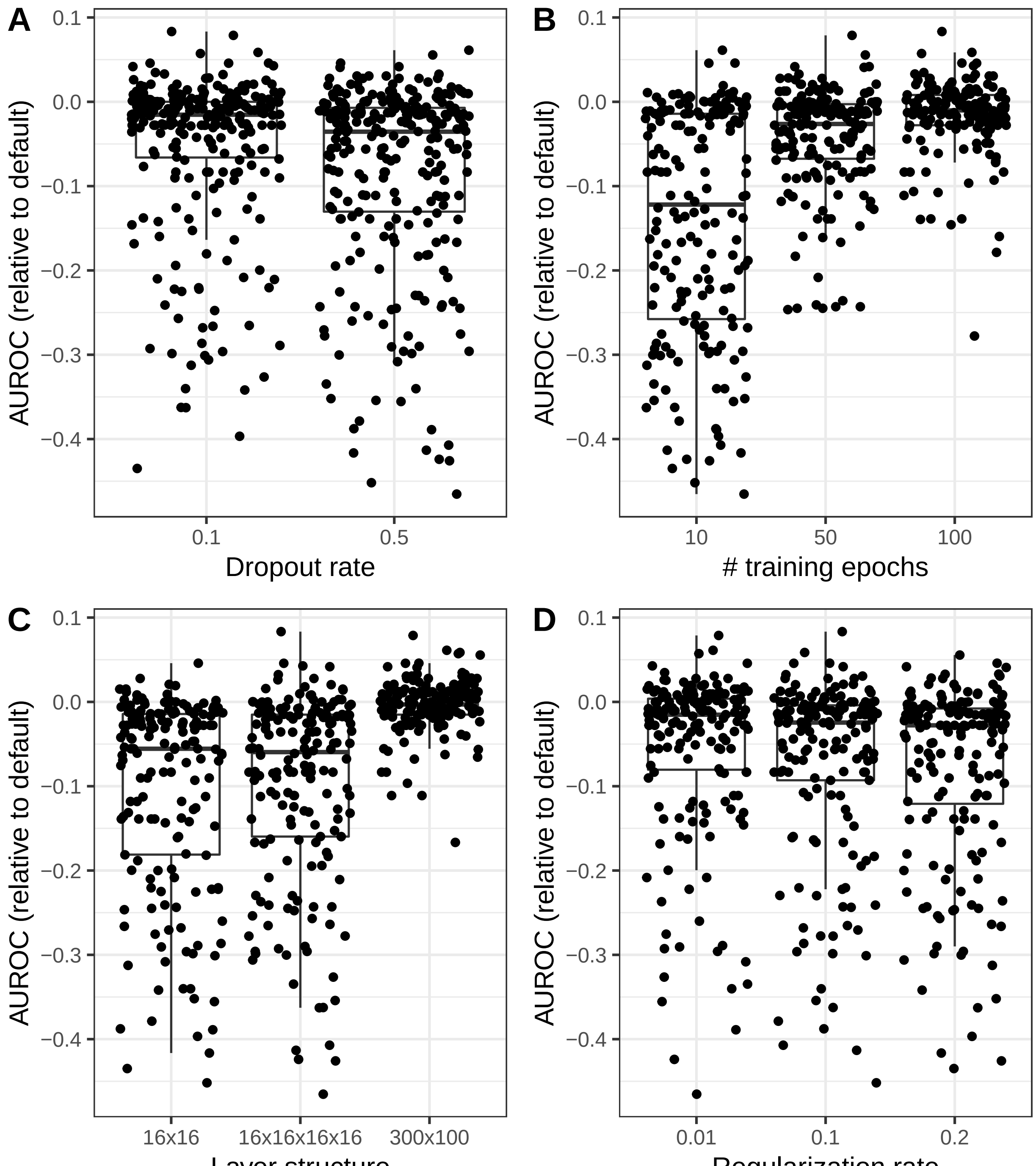


**Figure S9: Effect of changing different hyperparameters on performance of the keras/dnn algorithm.** We evaluated predictive performance for the `keras/dnn` algorithm using 53 different hyperparameter combinations. The level of performance varied depending on the hyperparameter combination used. Generally, AUROC values increased when using a smaller dropout rate, more training epochs, a wider layer structure, and a smaller regularization rate.

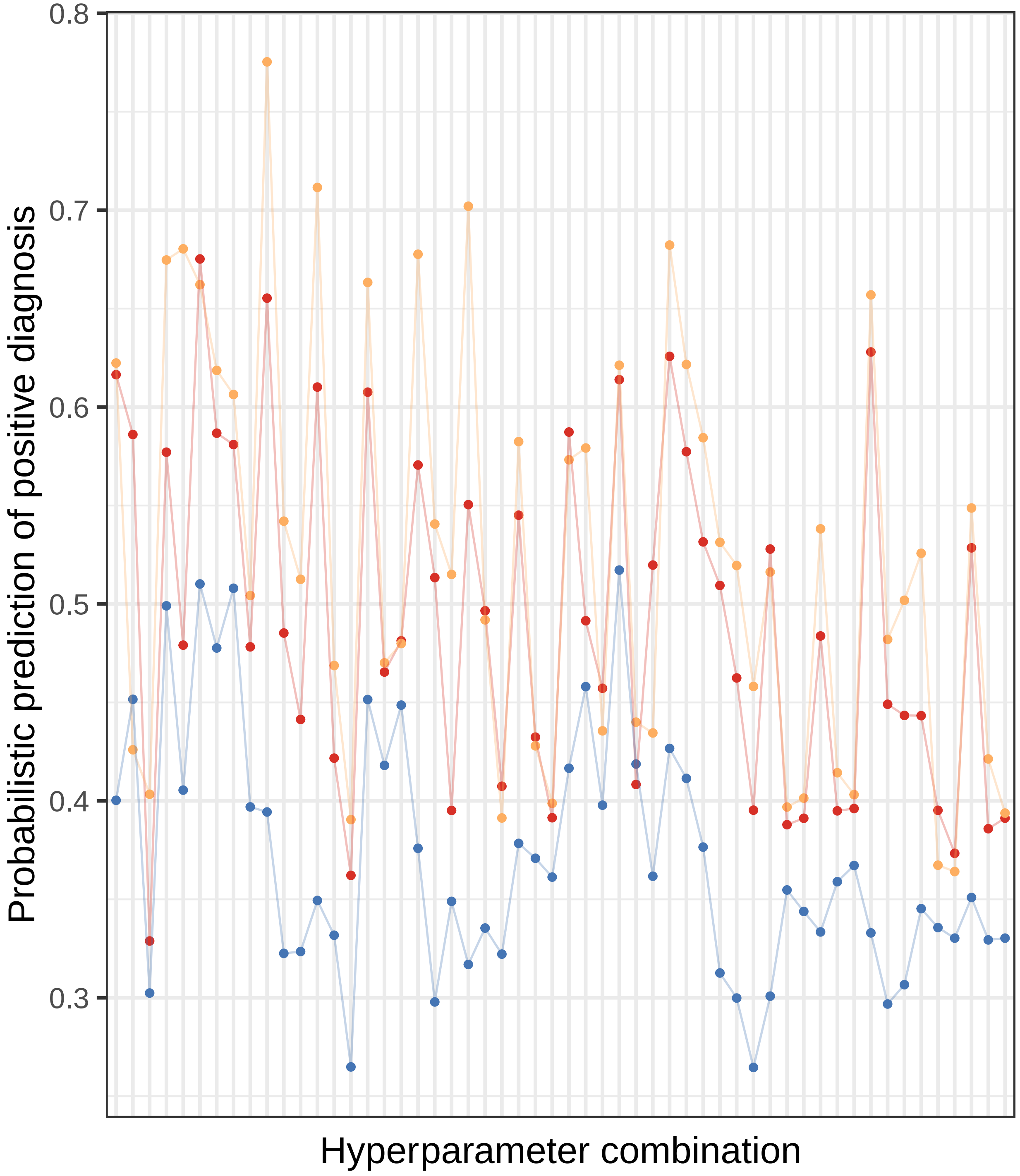


**Figure S10: Probabilistic predictions of positive diagnosis for patients in the Diabetes dataset for different hyperparameter combinations.** The Diabetes dataset includes a class variable indicating whether or not patients received a positive diagnosis. This figure shows probabilistic predictions of a positive diagnosis for three diabetes patients; the predictions were made using the `keras/dnn` algorithm. Each line represents predictions probabilities across different hyperparameter combinations.

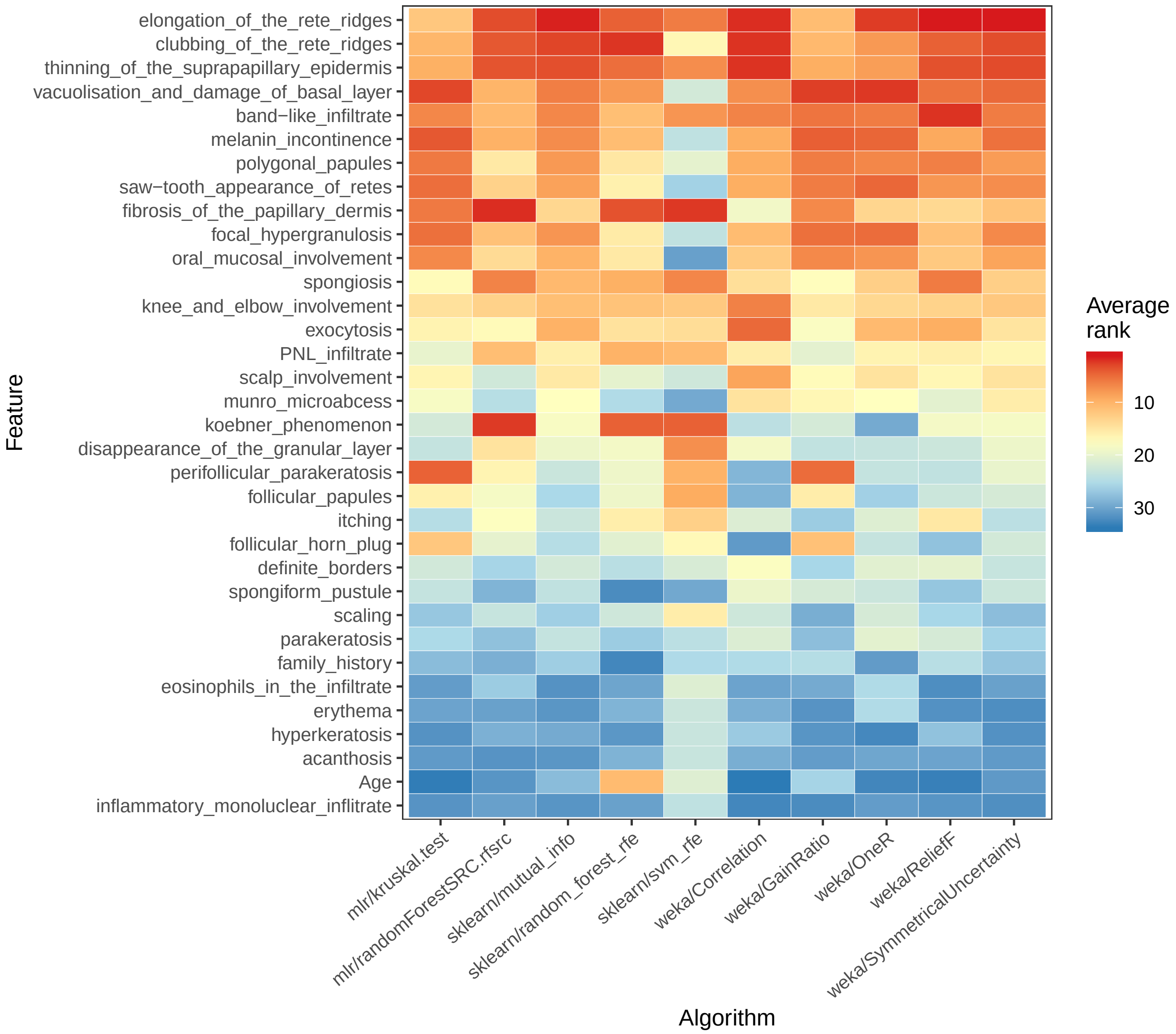


**Figure S11: Rankings of each feature in the Dermatology dataset for each feature-selection algorithm.** In this example, 10 feature-selection algorithms were applied to the Dermatology dataset. Each cell represents the average rank of each feature across nested cross-validation folds. Lower average ranks indicate greater relevance of the feature to the class variable (the patient's type of Eryhemato-Squamous disease). The average ranks were largely consistent across the feature-selection algorithms.
